# Autoantigen-specific CD8+ T-cell signature in Rheumatoid Arthritis

**DOI:** 10.64898/2026.02.18.706603

**Authors:** Janine Kemming, Helene Tenstad, Kristoffer Haurum Johansen, Kamilla Kjærgaard Munk, Birkir Reynisson, Christina H Bruvik Ruhlmann, Christian Nielsen, Søren Andreas Just, Sine Reker Hadrup

## Abstract

Recent evidence suggests that CD8+ T cells contribute to rheumatoid arthritis (RA) pathogenesis, however, their landscape of immune recognition, clonality, phenotypic and transcriptional characteristics, as well as functional properties remain poorly understood.

Using in silico epitope prediction, barcoded pMHC multimer screening, single-cell transcriptomic and TCR repertoire analysis in blood and tissue of RA patients, we systematically interrogated CD8+ T-cell responses targeting RA-associated peptides.

First, we identified HLA class I-restricted CD8+ T-cell responses against RA autoantigens. Second, we demonstrate that epitope-specific CD8+ T cells from RA patients display a distinct transcriptional footprint compared to healthy donors, which is characterized by features consistent with antigen experience, cytotoxicity and effector differentiation. In parallel, our data indicate that citrullination can modulate cross-recognition between peptide epitopes, suggesting that this post-translational modification may broaden or reshape antigen recognition within the CD8+ T-cell repertoire. Finally, we show that RA-associated TCR clonotypes are stable in peripheral blood over time, are comprised of phenotypically antigen-experienced cells, and can be detected in synovial tissue.

Together, our study defines a set of HLA class I-restricted CD8+ T-cell epitopes associated with RA and provides mechanistic insight into how citrullinations may influence CD8+ T-cell recognition as well as interrogating the RA-associated CD8+ T cell clonotype landscape. These findings support a direct role for autoreactive CD8+ T cells in RA and provide a foundation for targeted immunotherapy.

## Introduction

Rheumatoid arthritis (RA) is a chronic, systemic autoimmune and autoinflammatory disease, affecting approximately 1% of the world’s population. RA is characterized primarily by inflammation and progressive destruction of bone and cartilage, but inflammation also occurs in connective tissues in other organs such as lungs, eyes and vessels. Suspected targets for RA autoreactivity are proteins commonly found in connective tissues, including synovium, such as Aggrecan, Collagen, Enolase, Fibrinogen, Grp78, and Vimentin. To date disease-modifying anti-rheumatic drugs (DMARDs) alleviate symptoms and slow disease progression, but no curative treatment exists. Despite advances in DMARD treatment, a subset of patients continues to experience highly active disease due to drug resistance or intolerable side effects, resulting in substantial impairment of quality of life (REF). Disease activity is commonly assessed using the Disease Activity Score in 28 joints with C-reactive protein (DAS28-CRP), a composite clinical index integrating tender and swollen joint counts, systemic inflammation measured by CRP, and patient global assessment to quantify disease severity and monitor treatment response over time.

Existing evidence demonstrates a central role for CD4+ T cells in the pathogenesis of RA^1^. This is supported by the strong genetic association of RA with specific major histocompatibility complex (MHC) class II alleles^2–4^ and by the clinical efficacy of blocking T-cell co-stimulation using CD80/CD86 inhibitors such as abatacept^5^. Together, these findings support a role for CD4+ T cells as drivers of disease initiation and progression.

One hallmark of RA is the infiltration of T cells into inflamed synovial joints, where chronic inflammation promotes the overexpression and activation of peptidyl-arginine deiminases (PADs)^6,7^ . PAD activity reshapes the local immunopeptidome by converting peptidyl-arginine into peptidyl-citrulline, generating neo-epitopes that are immunologically distinct from their native counterparts. In RA, citrullinated peptides are targeted by anti-citrullinated protein antibodies (ACPAs), which are widely used as diagnostic and prognostic biomarkers and correlate with disease severity and joint damage^8^. These observations underscore the central role of citrullination in linking CD4+ T-cell help to pathogenic B-cell responses.

Beyond CD4+ T cells, emerging evidence indicates that citrullinated antigens can also be recognized by CD8+ T cells^9^. Expanded, activated, and cytotoxic CD8+ T-cell clones have been identified in both synovial tissue and peripheral blood of RA patients^10–12^, and CD8+ T cells from RA patients display enhanced reactivity toward citrullinated, but not native, vimentin^9^. Moreover, human leukocyte antigen (HLA)-B*08 (Asp9) is a known, moderate genetic risk allele for RA^13^. Despite these converging lines of evidence implicating autoreactive CD8+ T cells in RA pathogenesis, fundamental aspects of this response remain poorly defined. In particular, the frequency, epitope specificity, phenotype, and clonality of RA-specific CD8+ T cells are largely unknown. Addressing these gaps is essential for a more complete understanding of RA immunopathology and may open new avenues for targeted immunotherapeutic interventions.

Thus, in this study we set out to comprehensively describe the RA-specific CD8+ T cell landscape of immune recognition using barcoded peptide-MHC (pMHC) libraries and utilize the identified epitopes to analyze native versus citrullinated immunity, differences between patients and healthy donors as well as dynamics of clonality in both peripheral blood and the site of inflammation, the synovial tissue.

## Results

### Landscape of CD8+ T cell immune recognition in rheumatoid arthritis

We in silico predicted a library of 424 HLA-binding peptides stemming from RA-associated proteins collagen, vimentin, fibrinogen, enolase, GRP78 and aggrecan, restricted by the HLA alleles HLA-A*01:01, A*02:01, A*24:02, B*08:01, C*07:01 and C*07:02 (Table S1). If a candidate epitope contained one or more arginine residues, a second in silico prediction round was performed in which these residues were replaced by an undefined amino acid, and peptides that retained predicted HLA binding were included in the peptide library. This library, together with a library of 276 viral peptides, was tested for T cell recognition in a cohort of 40 HLA-matched RA patients (20 late RA, 20 early RA) as well as 17 healthy donors (HDs) using a DNA-barcoded pMHC-multimer screening technology (Fig.1A/B, Table S2, FigS1F). “Late RA” identified patients with long-standing disease diagnosis and pre-treatment while “early RA” patients were recently diagnosed and remained untreated at timepoint 1. Blood samples were collected over 3 different timepoints, each 3 months apart, while HDs were only sampled on timepoint 1 (Fig 1A). We sorted virus and RA-specific CD8+ T cell populations (Fig 1C) and deconvoluted barcoding data with barracoda 2.0 (DTU). Figure 1D shows the landscape of CD8+ T cell recognition for time point 1 (see Fig S1A/B for timepoint 2/3), with citrullinated epitopes accounting for more than half of the T cell responses detected (Fig S1C-E). Overall, we could detect T cell recognition towards 48 distinct epitopes and a total of 316 T cell responses targeting these epitopes. The protein distribution of recognized epitopes mirrored the initial library (Figure 1E/F) and responses were detected for all tested HLA restrictions. 9/48 epitopes had a prevalence of T cell recognition exceeding 40% of tested HLA-matched donors, whilst for the majority of epitopes, T cell recognition was rare (median 10% prevalence) (Fig 1G).

**Figure 1.**
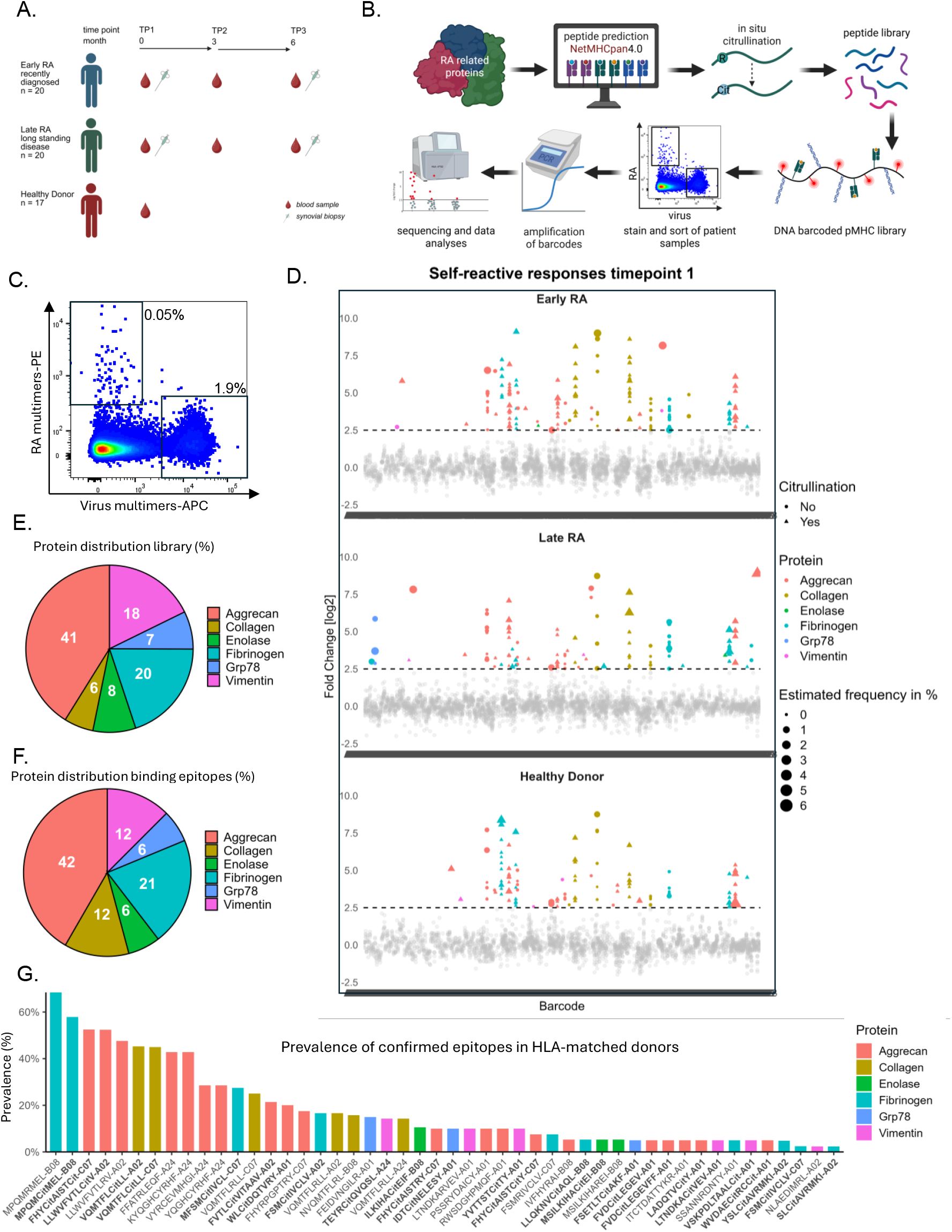
Barcode-labeled pMHC screen for epitope identification. A) PBMCs and synovial biopsies from 20 Patients with recently diagnosed rheumatoid arthritis (earlyRA), 20 patients with long standing RA (late RA) and 17 HD were obtained at the timepoints indicated. B) Schematic of the workflow, RA-specific CD8+ T cell epitope candidates were in silico predicted from RA-relevant proteins and in silico citrullinated. Barcode-labeled pMHC multimers were used to screen for responses against epitope candidates. C) Representative FACS plot of a patient sample stained with RA and Virus multimers. D) Dot plot showing significant (colored) CD8+ T cell responses against RA epitope candidates at timepoint 1 (triangles = citrullinated peptides, circles = native peptides). Colors refer to proteins and size of the dot indicates the estimated frequency. E) Protein distribution within the library and the confirmed CD8+ T cell epitopes (F). G) Prevalence of all identifies CD8+ T cell epitopes within the entire cohort. Citrullinated epitopes in bold.

### Repertoire composition and temporal evolution of RA-specific CD8+ T cells

To investigate cohort-specific differences beyond the overall landscape described in Figure 1, we next compared the epitope repertoire breadth and longitudinal dynamics of RA-specific CD8+ T cells between HD, early RA, and late RA. HLA composition was balanced across cohorts (Fig. S1F), allowing direct comparison of epitope recognition metrics. Interestingly, the number of recognized epitopes per individual did not significantly differ between HD and patients across timepoints. (Fig. 2A). Despite similar per-individual epitope counts, repertoire composition differed markedly at the cohort level. While HD responses were largely restricted to a limited set of shared epitopes, patients collectively recognized a substantially broader spectrum of RA-associated epitopes (Fig. 2B). Specifically, 65% of detected epitopes were uniquely observed in patients, whereas only 4% were uniquely detected in HD (Fig. 2B). We next examined the longitudinal behavior of individual epitope-specific CD8+ T-cell frequencies (Fig. 2C). Overall, frequencies in HD and patients were within a comparable range. However, early RA patients showed a trend toward increased epitope-specific frequencies at TP3 compared with TP1 (p = 0.07) (Fig. 2C, middle panel), whereas no clear directional change was observed in late RA. To assess the relationship between RA-specific CD8+ T-cell magnitude and clinical disease activity, we correlated the summed RA-specific T-cell frequency with DAS28 (Fig. 2D/E). In early RA, there was a tendency toward increased cumulative RA-specific T-cell frequency at TP3 despite decreasing DAS28 scores, reflecting the trend observed in Fig. 2C. In contrast, late RA patients demonstrated a significant positive correlation between summed RA-specific CD8+ T-cell frequency and DAS28 (rmcorr r = 0.54, p = 0.036; Fig. 2E). In this cohort, decreasing disease activity was associated with reduced RA-specific CD8+ T-cell frequencies, indicating that autoreactive CD8+ T-cell magnitude might more closely track clinical inflammation in established disease.

**Figure 2.**
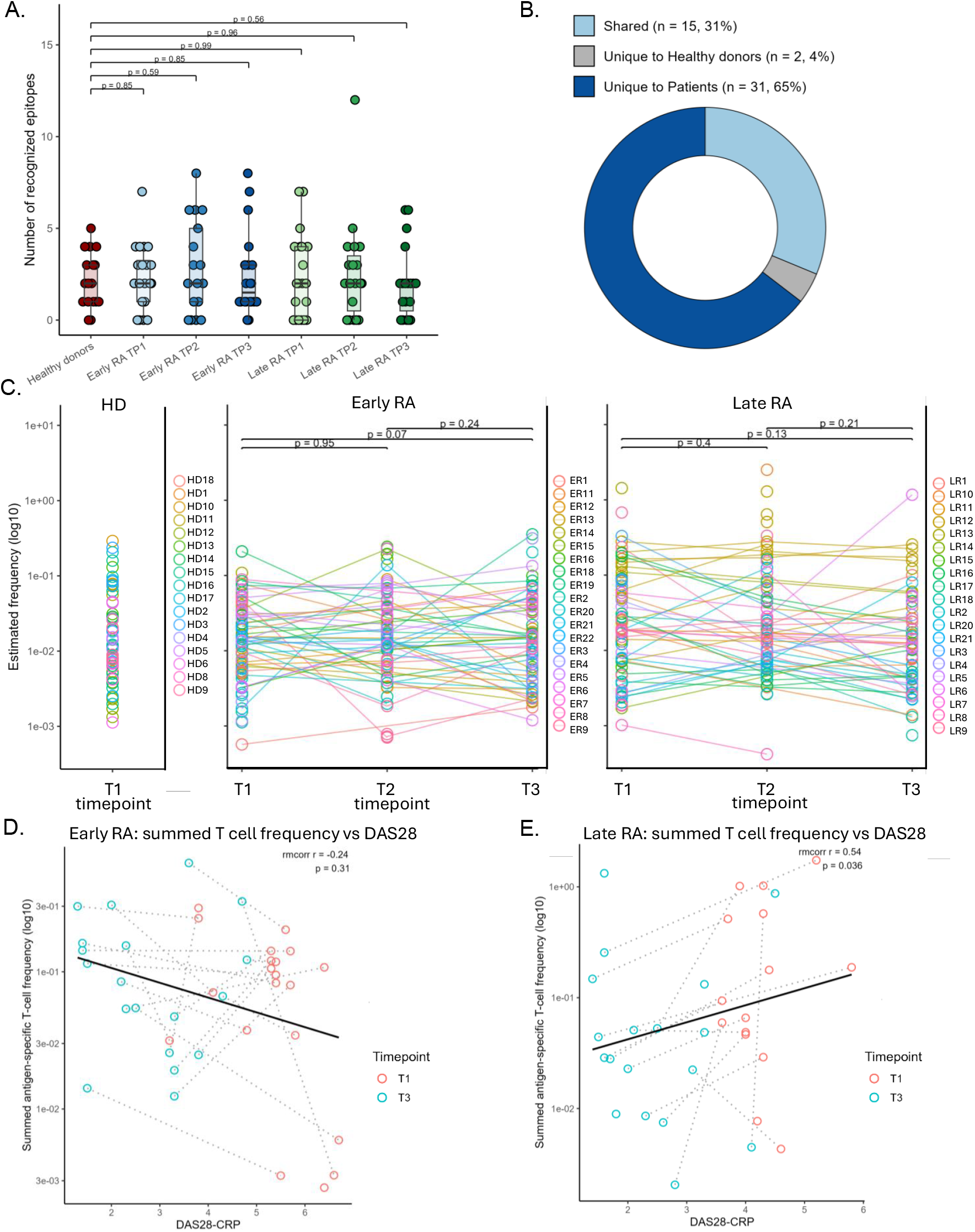
Repertoire composition and temporal evolution of RA-specific CD8+ T cells. A) Number of recognized RA-associated epitopes per individual in healthy donors (HD), early RA (TP1-TP3), and late RA (TP1-TP3). Dots represent individuals; boxes indicate median and interquartile range. Group comparisons were performed using two-sided Wilcoxon rank-sum tests. B) Cohort-level epitope distribution. Donut chart shows epitopes shared between cohorts or uniquely observed in HD or patients. Epitopes were defined using stringent criteria (logFC > 2.5, p < 0.001, estimated frequency > 0.01%, count ≥ 10) C) Longitudinal evolution of individual epitope-specific CD8+ T-cell frequencies (log10 scale). Lines represent patient×epitope trajectories. Paired comparisons between timepoints were performed using two-sided Wilcoxon signed-rank tests (LR=LateRA, ER =EarlyRA). D-E) Correlation between summed RA-specific CD8+ T-cell frequency and DAS28-CRP in early RA (D) and late RA (E). Points represent individuals at TP1 and TP3; black line indicates linear regression. Within-individual associations were assessed using repeated-measures correlation (rmcorr). All tests were two-sided; p-values are shown.

### RA-specific CD8+ T cell signature in patients

As the pMHC barcoding screen revealed that RA-specific CD8+ T cells could be detected both in patients and HDs, we set out to decipher differences between these cells in patients versus HDs by analyzing self-epitope specific CD8+T cells on the single cell level. We selected a cohort of 3 early RA, 5 late RA patients and 3 HD (Table S2) and analyzed the 12 most frequently detected RA and 21 most frequently detected viral epitopes from the pMHC barcoding screen (Fig 1G, Fig 3A,Table S1). Single-cell transcriptomes were integrated using Seurat and visualized by uniform manifold approximation and projection (UMAP). The resulting transcriptional landscape resolved 9 major clusters (Fig 3B) corresponding to distinct differentiation and lineage states as revealed by cell type annotation with ProjecTIL^14^ (Fig 3C). Integration with pMHC multimer barcode data using iTRAP2^15^ (Fig S2) linked transcriptional clusters to their cognate antigen specificities. If iTRAP2 assigned 2 specificities with equal confidence and significance, the respective TCRs were considered cross-binding to both pMHC multimers. This cross-reactivity was observed for an HLA-A*24:02 RA peptide epitope length variant, an HLA-B*08:01 citrullinated/native RA peptide pair and interestingly, HLA-C*07:02 restricted epitope MFSMCitIVCL and cytomegalovirus (CMV)-specific epitope CRVLCCYVL (Fig S3). Distinct pMHC combinations mapped to discrete subpopulations within the UMAP revealing antigen-driven heterogeneity (Fig 3D). Interestingly, the HLA-C*07:02-restricted RA/CMV-specific CD8+ T cells formed a prominent subset in all HLA-C*07:02+ donors, clustered away from HLA-A/B RA-specific and virus-specific CD8+ T cells and were thus analyzed separately (referred to as RA HLA-C). RA-specific HLA-A/B-restricted CD8+ T cells (RA HLA-A/B) tended to contain more naïve cells than virus-specific (Virus) and RA HLA-C CD8+ T cells (Fig S3D) which seemed to be driven by a prominent naïve signature in the HDs compared to patients (Fig 3E, Fig S3A-D), albeit non-significant due to small sample size. Differential gene expression (DEG) analysis across pMHC categories RA HLA-C, RA HLA-A/B and Virus (Table S1) revealed transcriptional programs distinguishing the cell subsets. A heatmap of top cluster markers (adjusted p < 0.05) illustrated most significantly upregulated transcripts in the RA HLA-C subset, with upregulation of adaptive, Natural Killer (NK)-like signatures including Killer Immunoglobulin-like receptors (KIRs) KIR2DL3, KIR3DL2, KIR3DL1 and KIR2DL1, Killer Cell Lectin Like Receptor C2 (KLRC2, also: NKG2C), tyrosine kinase-binding protein (TYROBP), Killer Cell Lectin Like Receptor F1(KLRF1), Helios (IKZF2) as well as drivers of cytotoxicity and antigen-recognition; granzyme B (GZMB), Cathepsin W (CTSW) and Acyloxyacyl Hydrolase (AOAH) (Fig3F). The RA HLA-A/B signature confirmed a more naïve-like/ central memory phenotype driven by SELL (CD62L) expression (Fig3F/G) as well as RA associated genes like Coactosin Like F-Actin Binding Protein 1 (COTL1) (Fig3F/H). To assess disease associated remodeling within the RA HLA-A/B CD8+ T cell subset, we compared gene expression profiles of these cells between patient groups and healthy donors revealing a clear patient-specific cytotoxic (GZMA, GZMB, perforin (PRF1)), activating and migration (CC Motif Chemokine Ligand 5 (CCL5), CD81, Integrin Subunit Beta 2(ITGB2)) associated transcriptional program (Fig 3I-K). Upregulation of HLA-A/C expression in patients’ CD8+ T cells was consistent with an inflammatory environment and gene modules like Acyloxyacyl hydrolase (AOAH) and Cathepsin C (CTSC) supported a transcriptional program associated with enhanced inflammatory responsiveness and cytotoxic effector function. In contrast HD-derived counterparts with shared pMHC specificity preferentially expressed genes associated with homeostatic signaling (Regulator of cell cycle (RGCC), Lymphoid enhancer-binding factor 1(LEF1), Lymphotoxin-beta(LTB), cytokine responsiveness (Leptin Receptor Overlapping Transcript Like 1(LEPROTL1), Fms Related Receptor Tyrosine Kinase 3 Ligand (FLT3LG)) and quiescent memory states (Fig 3I). Gene Ontology (GO) enrichment confirmed strong signatures of cell killing, inflammatory response and cytotoxicity only in RA HLA-A/B CD8+ T cells of patients (Fig 3L). Differential expression analysis of RA HLA-C CD8+ T cells revealed distinct transcriptional programs between patient and HD cells (Fig 3M). Patient-derived RA HLA-C CD8+ T cells showed upregulation of NK-like markers (KIRs, KLRB1) (Fig 3N/O) as well as cytotoxic effector molecule GZMM (Fig 3M) while HD derived counterparts displayed distinct transcriptional programs of (effector) memory T cells with expression of transcription factor Hobit (ZNF683), granulysin (GNLY) and LEF1. GO enrichment confirmed several cytotoxicity-related pathways significantly overrepresented in patient-derived compared to HD counterparts within the same RA HLA-C specific CD8+ T-cell population (Fig. 3P). Interestingly, T cell binding of HLA-C*07:02 restricted MFSMCitIVCL/ CRVLCCYVL multimers seemed to be at least partly due to binding to KIR instead of TCR, as demonstrated through blocking of KIR, abolishing pMHC multimer binding (Fig S3E).

**Figure 3.**
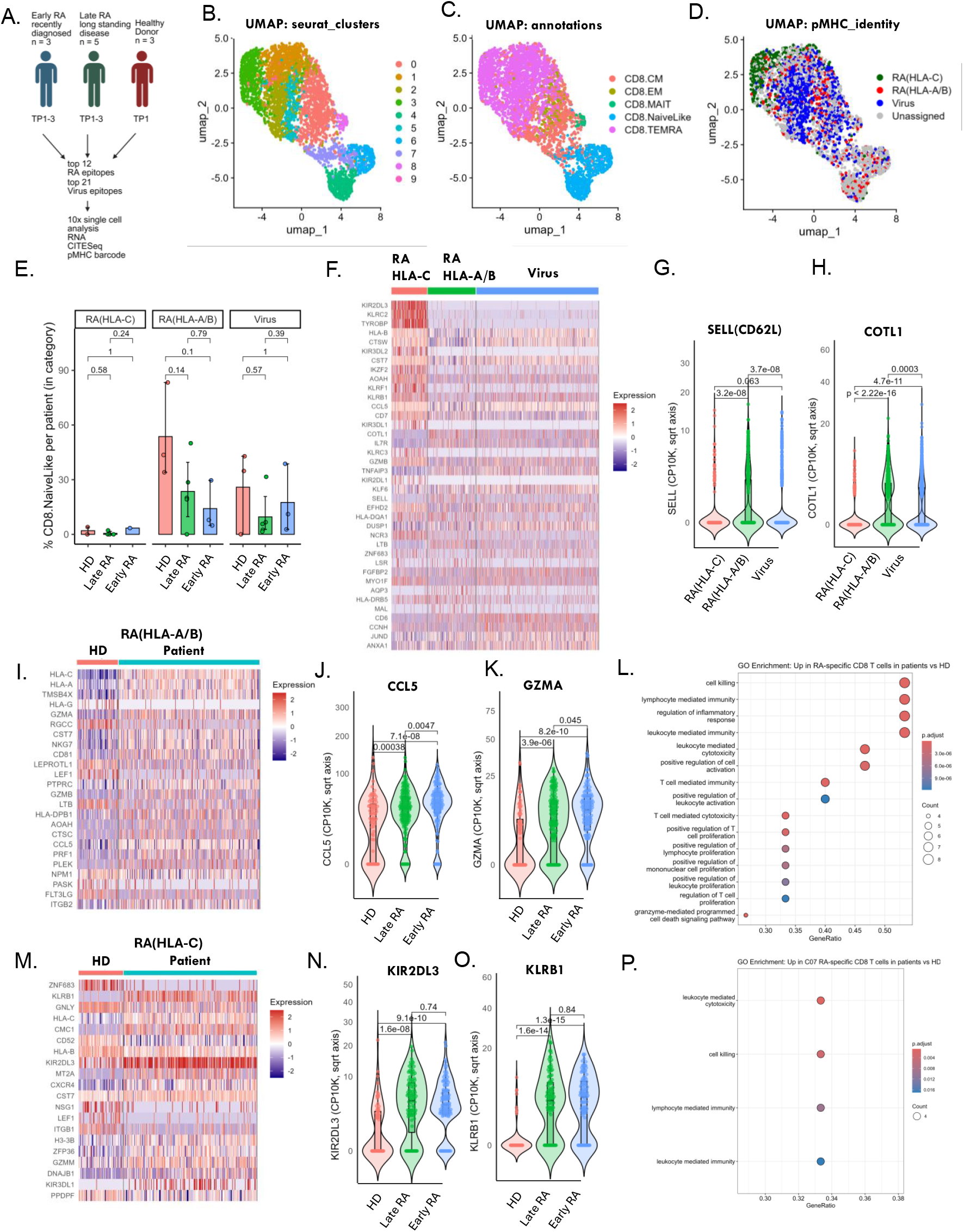
single-cell analyses of RA-specific CD8+ T cells. A) Schematic of patient cohort, library and workflow for the 10x 5’ single cell experiment. B) UMAP of all cells colored by Seurat-derived unsupervised clusters, C) projected cell type annotations (ProjecTIL) and D) pMHC–TCR barcode assignments (RA-specific, HLA-A/B restricted: red; RA-specific. HLA-C restricted: green; Virus-specific: blue; Unassigned: grey). E) Barplot depicting the percentage of Naive-like cells in the different categories and patient cohorts. (F) Heatmap visualization of DEGs with adjusted p < 0.05 between categories. G-H) Expression values of IL7R and SELL as CP10K between categories. I) Heatmap visualization of DEGs with adjusted p < 0.05 in HLA-A/B restricted, RA-specific CD8+ T cells in patients vs HDs. J-K) Expression values of CCL5 and GZMA as CP10K. L) GO Enrichment showing pathways up in RA-specific CD8+ T cells of patients vs HDs. M) Heatmap visualization of DEGs with adjusted p < 0.05 in HLA-C restricted, RA-specific CD8+ T cells in patients vs HDs. N-O) Expression values of KIR2DL3 and KLRB1 as CP10K. and P) GO Enrichment showing pathways up in HLA-C restricted RA-specific CD8+ T cells in patients vs HDs

### Differences between citrullinated and native RA-specific CD8+ T cell epitopes

To dissect how citrullination alters antigen recognition by RA-specific CD8+ T cells, we compared native and citrullinated pMHC pairs restricted by HLA-A*02 and HLA-B*08. Across the pMHC-barcode screen, A02+ RA patients showed distinct and often divergent responses to the native versus citrullinated HLA-A*02:01 restricted epitope LLWVFVTLR/CitV (Fig 4A). Structural modelling indicated that the citrullinated residue protrudes from the pMHC complex and is positioned for direct contact with the TCR (Fig 4B). Consistent with this geometry, tetramer staining resolved two clearly separated T-cell populations in patients, indicating that LLWVFVTLRV and LLWVFVTLCitV are recognized by distinct T cell subsets (Fig 4C). The same pattern was observed for the native/citrullinated peptide pair VQMTFLR/CitLL (FigS4). In contrast, for the B08-restricted peptide pair MPQMR/CitMEL, the log fold change from the pMHC barcoding screen is identical for the citrullinated and native epitope in all B08+ donors (Fig 4D). Structural modelling revealed the citrullinated residue is oriented inward toward the HLA groove (Fig 4E) and patient tetramer staining showed a low affinity double-positive population in addition to some single positive cells suggesting that clones may have the potential to cross-recognize both native and citrullinated MPQMRMEL (Fig 4F). Transcriptomic profiling of B08-restricted T cells further supported this functional divergence: native-only, citrullinated-only, and cross-reactive (double-barcode) clonotypes exhibited distinct transcriptional signatures (Fig 4G). Notably, many cytotoxicity genes like GNLY (Fig 4G/H), Granzymes A/B/H and Perforin (Fig 4G) were uniquely upregulated in the cross-reactive CD8+ T cell subset and GO enrichment analysis highlighted the upregulation of cytotoxicity-associated pathways (Fig 4I). Together, these data indicate that the structural orientation of a citrullinated residue may dictate whether CD8+ T cells specific for the native epitope have the potential to cross-recognize the citrullinated version.

**Figure 4.**
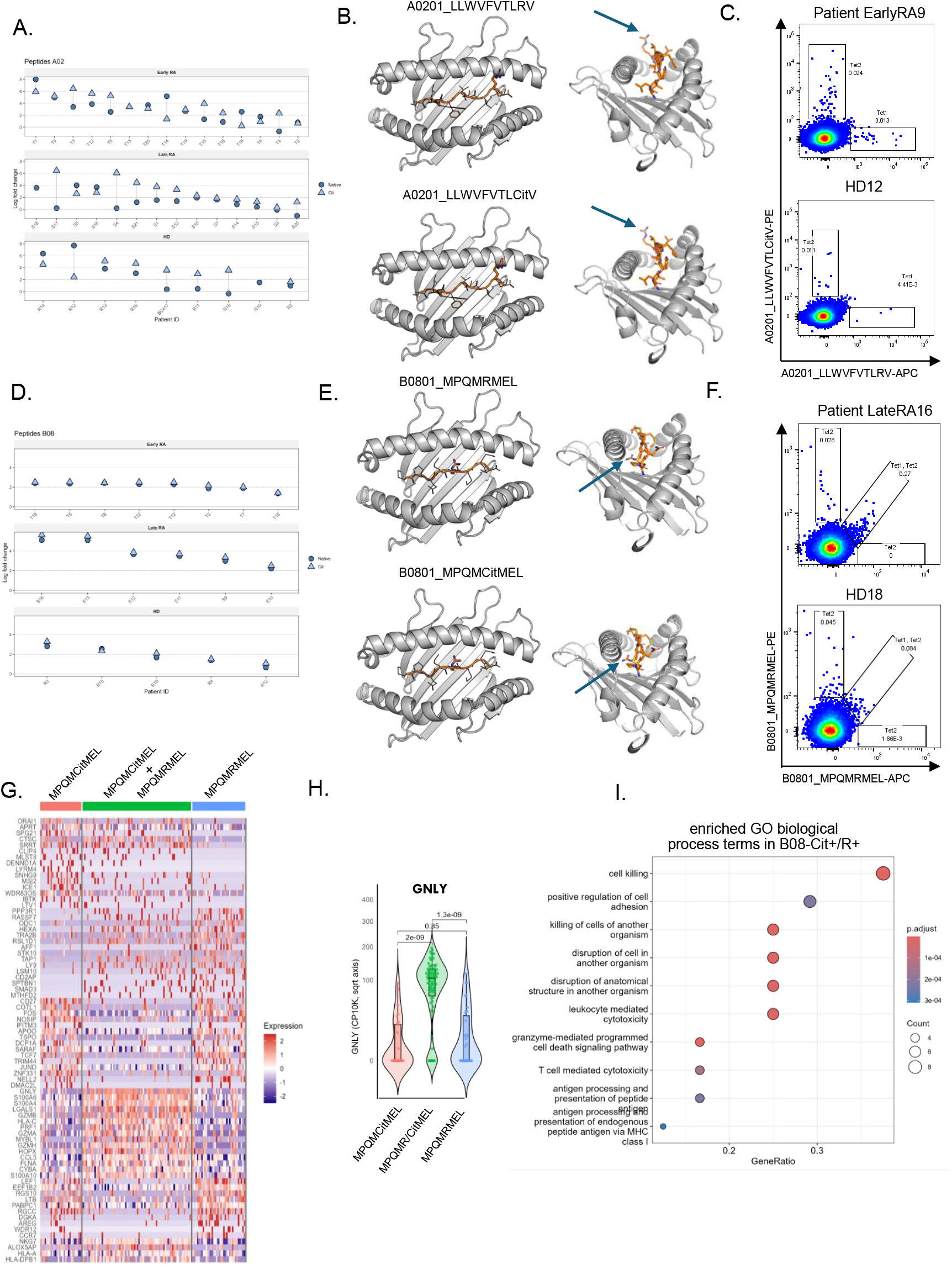
citrullinated vs native RA-specific CD8+ T cell epitopes. A) pMHC barcode screen data showing responses of HLA-A*02+ patients against RA-specific cit/native A*02-restricted peptide pair. B) Ribbon diagram of the HLA-A*02 monomer (grey) with the bound cit/native peptide ligand depicted (arrow indicates site of citrullination). C) Example FACS plot of tetramer staining with the cit/native peptide pair in patient vs HD. D) pMHC barcode screen data showing responses of HLA-B*08+ patients against RA-specific cit/native B*08-restricted peptide pair. E) Ribbon diagram of the HLA-B*08 monomer (grey) with the bound cit/native peptide ligand depicted (arrow indicates site of citrullination). F) Example FACS plot of tetramer staining with the cit/native peptide pair in patient vs HD. G) Heatmap visualization of DEGs for B*08 restricted cit/native and crossreactive RA-specific CD8+ T cells H) Expression values of GNLY as CP10K. I) Dot plot of enriched GO biological process terms, where dot size reflects the number of genes and color indicates adjusted p-value. Dot plot depicts the pathways enriched for CD8+ T cells crossreactive to the cit /native B∗08 restricted peptide pair vs the single epitopes

### Persistent peripheral TCR clones overlap with synovial T cells in RA

To define how circulating T cells relate to the synovial immune repertoire in RA, we integrated longitudinal PBMC single-cell TCR data with TCR sequencing from matched synovial biopsies. A schematic outlines the workflow from synovial biopsy to targeted TCR sequencing and data integration (Fig 5A). In representative patient EarlyRA19, the PBMC clonotype repertoire was remarkably stable across three longitudinal timepoints: dominant TCRαβ clonotypes persisted with similar relative abundance, demonstrating a conserved circulating T-cell landscape (Fig 5B). We next compared PBMC clonotypes from timepoint 1 to the patient’s timepoint 1 biopsy and observed clear overlap between compartments, indicating that the most expanded circulating T-cell clones are also present within the inflamed synovium, and a large fraction of these recognized RA-derived self-epitopes. A second patient LateRA20, analysed across PBMC timepoints 1 and 2, showed similar stability and likewise exhibited shared clonotypes between blood and synovial tissue (Fig 5C/D). The same pattern was observed for the remaining patients (FigS5A-H).

**Figure 5.**
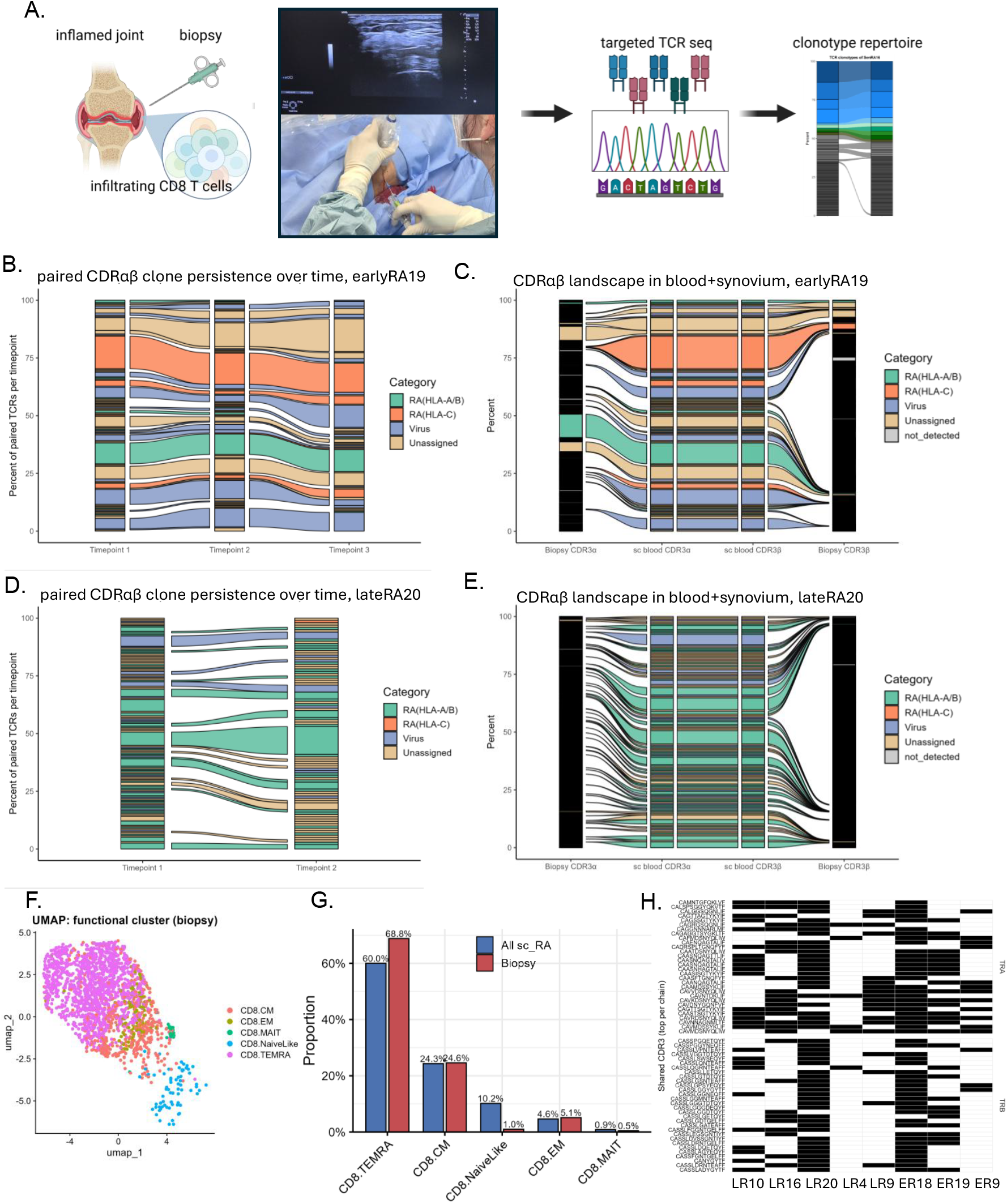
TCR clonotype landscape in blood vs synovial biopsies. A) Schematic visualizing TCR sequencing from synovial biopsies . B) Alluvial plot showing the persistence and dynamics of TCR clonotypes (paired alpha and beta) in a selected patient across timepoints 1-3. The width of each stream represents the relative frequency of a clonotype, colors correspond to category.C) Alluvial representation of TCR clonotypes shared between blood and synovial tissue compartments. D) Alluvial plot showing the persistence and dynamics of TCR clonotypes (paired alpha and beta) in a selected patient across timepoints 1-2. E) Alluvial representation of TCR clonotypes shared between blood and synovial tissue compartments. F) UMAP of the single-cell dataset showing cells carrying TCR a/β clonotypes also identified in biopsy repertoires, indicating overlap between peripheral and tissue T cell populations G) Proportional distribution of functional clusters among all annotated single-cell populations versus clonotypes overlapping with biopsy repertoires. H) Heatmap showing shared CDR3 sequences across patients identified in bulk synovial TCR sequencing, with TCRα chains displayed in the upper panel and TCRβ chains in the lower panel; black tiles indicate presence of the indicated CDR3 clonotype in a given patient (LR = LateRA, ER = EarlyRA).

Projecting biopsy-overlapping clonotypes onto the PBMC single-cell UMAP confirmed that synovium-associated TCRs corresponded to defined transcriptional states in circulation. Notably, these overlapping clonotypes were enriched in non-naïve T-cell clusters, including activated and cytotoxic subsets, whereas naïve cells contributed minimally (Fig 5F/G). These findings demonstrate that RA synovial tissue contained T-cell clones that are detectable and longitudinally stable in the blood, and that these shared clonotypes predominantly represent antigen-experienced effector populations rather than naïve T cells. Strikingly, bulk TCR sequencing of synovial biopsies revealed extensive sharing of identical CDR3α and CDR3β sequences across patients, however, due to the nature of bulk sequencing, α/β chain pairing could not be resolved (Fig. 5H). To assess functional reactivity, selected RA HLA-C specific TCRs were re-expressed by CRISPR-Cas9 knock-in and evaluated in an activation-induced marker (AIM) assay using peptide-pulsed, CD3-depleted primary target cells. Upregulation of CD137 and CD69 was only observed upon stimulation with the cognate peptide and not with irrelevant peptide (Fig. S6).

## Discussion

Previous studies demonstrated HLA class I-restricted CD8+ T-cell reactivity against citrullinated autoantigens in RA (Moon et al.), but minimal epitope-level resolution was lacking. Here, we define discrete HLA class I-restricted epitopes associated with RA and show how citrullination modulates their recognition. Our data support a direct pathogenic role for antigen-specific CD8+ T cells in synovial inflammation, rather than viewing them as passive bystanders. A major strength of this study is the convergence of independent layers of evidence. Epitope-specific CD8+ T-cell responses were first identified in a high-throughput pMHC barcode screen, reappeared in single-cell TCR datasets among transcriptionally defined effector populations, and were subsequently detected as expanded clonotypes in unbiased bulk sequencing of synovial tissue. The recurrence of identical TCR clonotypes across blood and inflamed joint strongly supports their disease relevance. Although pairing α/β chains in synovial bulk sequencing required integration with matched single-cell PBMC data, the reproducible detection of these antigen-specific clones across compartments argues that they represent genuine disease-associated clones.

The HLA-B*08:01-restricted fibrinogen α-chain epitope MPQMRMEL showed the highest prevalence. Its restriction by an established RA risk allele and detection in both native and citrullinated forms suggest biological relevance. Independent immunopeptidomics datasets from human synovium (SysteMHC Atlas^16^) confirm that the parent protein undergoes proteolytic processing at the site of pathology, supporting the plausibility of in vivo peptide liberation. Definitive validation of class I presentation in RA synovium will require direct immunopeptidomic mapping. Importantly, RA was not characterized by an increased number of epitopes recognized per individual, but by diversification of antigenic targets at the cohort level. Healthy donors predominantly targeted a restricted and overlapping epitope set, whereas patients collectively engaged a broader repertoire. This pattern argues against simple quantitative amplification and instead supports qualitative reshaping of antigenic selection. Although patients were sampled longitudinally at three timepoints versus one in healthy donors, per-individual breadth remained stable over time, indicating that diversification is unlikely to reflect sampling depth alone.

Notably, RA-associated epitope-specific CD8+ T cells were detectable in healthy donors, demonstrating that autoreactive specificities are present within the physiological repertoire. However, single-cell analyses revealed striking functional divergence: identical antigen specificities displayed predominantly naïve-like signatures in healthy donors, but cytotoxic and inflammatory programs in patients. These findings suggest that RA arises not from the de novo generation of autoreactive clones, but from functional reprogramming and expansion of a pre-existing autoreactive pool. The identification of defined class I epitopes provides a molecular foundation for precision therapeutic approaches. Strategies such as epitope-specific tolerogenic nanoparticles, targeted depletion, or engineered pMHC-based “minibinders”^17^ that block or delete pathogenic CD8+ T cell clones, tolerogenic pMHC nanoparticles, epitope-specific depletion strategies, and rational vaccine-based tolerance induction have remained out of reach in RA due to the absence of validated CD8+ T cell target epitopes. By pinpointing exact peptide specificities, our study establishes a framework for selective modulation of disease-driving CD8+ T cells. Our structural analyses further illustrate how citrullination can differentially influence TCR recognition in an allele-dependent manner. Although based on a limited number of peptide pairs, these observations highlight how residue orientation and MHC anchoring can shape neo-epitope formation versus cross-reactivity. Broader structural and biophysical analyses will be required to generalize these principles.

A prominent feature of our dataset was the transcriptional divergence between HLA-A/B and HLA-C-restricted RA-specific CD8+ T cells. HLA-C-restricted populations were dominated by an NK-like, KIR-associated cytotoxic program and formed a distinct cluster across donors. Although multimer binding within this compartment may partially involve KIR engagement, these cells were clonally expanded, displayed strong cytotoxic signatures in patients, and exhibited patient-specific transcriptional remodeling. Together, these findings suggest that HLA-C-restricted RA-specific CD8+ T cells occupy a hybrid adaptive/innate state that is selectively skewed toward effector function in disease.

Longitudinal TCR analyses revealed remarkable stability of dominant peripheral clonotypes over time, consistent with previous reports of oligoclonal RA repertoires. By linking PBMC single-cell data to matched synovial repertoires, we show that circulating clones overlap with joint-infiltrating populations and are biased toward activated, cytotoxic clusters rather than naïve states. This supports a model of persistent, recirculating effector clones contributing to chronic inflammation.

In summary, we provide the first systematic definition of the HLA class I-restricted CD8+ T-cell epitope landscape in RA. By integrating large-scale pMHC screening, single-cell transcriptomics, TCR sequencing, and synovial repertoire analysis, we demonstrate that RA-specific CD8+ T cells are clonally expanded, functionally reprogrammed, and shared between blood and joint. Our data indicate that autoimmune pathology is characterized by diversification and expansion of a pre-existing autoreactive repertoire, coupled to functional activation and reshaping of the antigenic space engaged in disease. These findings establish a molecular framework for understanding CD8+ T-cell-mediated autoimmunity in RA and open avenues for epitope-specific precision immunotherapy.

## Material and Methods

### Study cohort

A total of 20 patients with long-standing RA (LateRA), 20 patients with recently diagnosed RA (EarlyRA) and 17 HLA and age-matched healthy donors (HD) were included. HD7, EarlyRA10/17 and LateRA19 were excluded from the original cohort due to an unmatching HLA profile (data not shown). All donors were recruited at Odense University Hospital. All donors were tested for autoantibodies and other relevant disease parameters (Table S2). Patients PBMC samples were obtained at 3 different timepoints, 3 months apart, whilst healthy donor samples were only collected at time point 1. Relevant patient characteristics can be found in Table S2. Next-generation sequencing was used for HLA typing. Written informed consent was obtained from all donors and the study was conducted according to ethics committee regulations (vote: 110272) and the Declaration of Helsinki.

### Synovial biopsies

At baseline and 6-month visits, synovial biopsies were obtained by USGSB in a clean procedure room, as previously described^18^. All biopsies were taken from the same wrist at baseline and 6 months. Briefly, local anaesthetic was injected into the soft tissue up to the joint capsule and into the joint space and a Quick-Core biopsy needle (16-gauge; Cook Medical) was then guided by ultrasound and placed within the joint capsule to retrieve synovium. The biopsies were taken over the scaphoid and lunate junction in the radiocarpal joint of the wrist.

### RA peptide prediction and selection

Candidate HLA class I-restricted peptides were predicted using NetMHCpan v4.0 (Technical University of Denmark, http://www.cbs.dtu.dk/services/NetMHCpan/). Protein sequences from six RA-associated autoantigens (aggrecan, collagen type II, enolase, fibrinogen, GRP78, and vimentin) were screened for 8–12-mer peptides predicted to bind HLA class I alleles HLA-A*01:01, HLA-A*02:01, HLA-A*24:02, HLA-B*08:01 and HLA-C*07:02 included in this study. Peptides with predicted binding ranks (%Rank) ≤ 1 were classified as binders.

To identify potential citrullination-dependent epitopes, an additional in silico prediction strategy was applied in which arginine residues were systematically substituted with a hypothetical amino acid (“X”) to remove arginine-specific anchor constraints. Peptide sequences that were not predicted to bind in their native form but achieved binder status following substitution were included as candidates representing potential citrullinated variants.

In total, 424 peptides predicted to bind one or two HLA molecules were selected for experimental validation (Supplementary Table 1). All predicted peptides, together with 276 viral control peptides, were synthesized by Pepscan Presto (The Netherlands) at a 2 µmol scale. Peptides were quality controlled by the manufacturer using UV spectroscopy and mass spectrometry (random 5% sampling) and dissolved in DMSO to a stock concentration of 10 mM. Sequences and details of the peptide libraries can be found in Table S1.

### Assembly of DNA barcode–labeled dextran multimer libraries

DNA barcode–labeled dextran multimer libraries were assembled using biotinylated combinatorial DNA barcodes^19^ (LGC Biosearch, 2.17 μM), fluorescent streptavidin– dextran conjugates (Fina Biosolutions Inc., 160 nM), and custom recombinant, biotinylated, UV-cleavable pMHC monomers (50 μg/ml, ∼1 μM). Fluorescent dextran conjugates were first centrifuged twice at 10,000g for 2 min at 4°C to eliminate aggregates, then preincubated with biotinylated DNA barcodes at 4°C for 30 min, using a molar ratio of 0.5 DNA barcodes per dextran. Peptide-loaded pMHC monomers were centrifuged separately (3300g, 5 min, 4°C), and the supernatant was then added to the barcoded dextran conjugates at a ratio of 16–18 pMHC monomers per dextran. To stabilize the complexes, a premixed freezing buffer was added, yielding final concentrations of phosphate-buffered saline (PBS), 1.5 μM d-biotin, 0.1 mg/ml Herring DNA, 0.5% bovine serum albumin, 2 mM EDTA, and 5% glycerol.

The assembled DNA barcode-labeled multimers were incubated for 20 min at 4°C before being stored at −20°C. The starting concentration of the dextran backbone was 160 nM, and the final concentration of the assembled multimer was 35.56 nM prior to staining.

### Barcoded pMHC multimer staining and amplicon sequencing

Per staining reaction 1.5 μl of every assembled DNA barcode–labeled multimer was pooled. For the RA library multimers were initially grouped into HLA-restricted pools and subsequently subdivided into 2-digit HLA-matching, patient-specific pools, the viral library was stained for each patient independent of HLA match. The multimer pools were upconcentrated and centrifuged twice to remove aggregates before use. Each sample was stained with the multimer pools for 30 min at RT, followed by labeling with surface antibodies (Table S3) and viability dyes at 4°C and fixation with 1% PFA in PBS. Fluorophore-positive, antigen-specific populations were sorted on a BD Aria Fusion fluorescence-activated cell sorter and spun down at 5000xg for 10 min at 4C. Co-attached DNA barcodes from sorted antigen-specific populations were subsequently amplified using Taq polymerase together with sample-indexed primers, as described previously^19^. The amplified PCR products were purified with the QIAquick PCR purification kit (QIAGEN) and sent for sequencing at PrimBio (USA). Resulting data was analyzed using Barracoda 2.0 (https://services.healthtech.dtu.dk/tuba/barracoda-2.0/) as previously described and responses were required to pass significance criteria (p <0.001, log2fold change >2.5, count1 >10) and exceed a minimum estimated frequency of 0.01% of the parent population to reduce low-count barcode noise.

### Single-cell sequencing

PBMCs were thawed, washed twice in Cell Staining Buffer (PBS + 0.5% BSA), and resuspended in 20 µL of buffer and a patient specific pMHC multimer panel containing sc barcodes was added and incubated for 60 min at 4°C, followed by 3x washing steps and resuspension in 20 µL Cell staining Buffer with 2.5 µL Human TruStain FcX™ Fc Blocking reagent (BioLegend, Cat#42302) for 10 min at 4°C. After 10 min reconstituted TotalSeq-C Human Universal Cocktail (BioLegend) was added together with a surface antibody panel (Table S3) and 0.5 µL of a sample-specific TotalSeq-C anti-human Hashtag Antibody (BioLegend). Cells were then incubated for 30 min at 4°C, washed 3x in Cell Staining Buffer and finally reconstituted in 150 µL Cell Staining Buffer and kept at 4°C until sorting.

Antigen-positive CD8+ T cells were identified based on PE/APC-labeled DNA-barcoded pMHC multimers and sorted using a FACS Melody (BD Biosciences). Approximately 25000 sorted cells were pooled into a single tube containing 100 µL of Cell Staining Buffer, centrifuged at 390 × g for 10 min at 4°C, and the supernatant discarded. The resulting stained cells were processed using the 10x Genomics platform with 5′ v2 chemistry, enabling combined use of immunophenotyping antibodies and DNA-barcoded pMHC multimers.

Downstream processing of mRNA and DNA barcodes was performed according to the manufacturer’s protocol (Chromium Next GEM Single Cell 5′ Reagent Kits v2, Dual Index; 10x Genomics, USA), incorporating Feature Barcode technology for Cell Surface Protein and Immune Receptor Mapping. TCR, barcode (pMHC multimers, immunophenotyping antibodies, and hashtag antibodies), and gene expression libraries were generated and sequenced by Novogene on a NovaSeq system (Illumina) using a 150 bp paired-end program.

### Construction of single-cell libraries

Single-cell libraries were generated with the 10x Genomics Chromium Next GEM Single Cell 5′ v2 chemistry, applying Feature Barcode technology for Cell Surface Protein and Immune Receptor Mapping, according to the manufacturer’s instructions (10x Genomics, USA). Sorted cells were loaded onto a Chromium Next GEM Chip and processed in a Chromium Controller to create Gel Beads-in-Emulsion (GEMs). cDNA was synthesized from mRNA, while DNA was derived from Feature Barcodes.

Targeted amplification was performed for 16 cycles, and products were size-separated using SPRIselect beads (Beckman Coulter, Cat#B23318). This generated three library types: TCR (VDJ), Gene Expression (GEX), and Barcode (BC). Libraries were quantified with the Qubit dsDNA HS Assay Kit (Invitrogen, Cat#Q32851), pooled at a ratio of 1 BC : 5 GEX : 1.5 TCR, and sequenced on an Illumina NovaSeq platform (Novogene, UK) with 150 bp paired-end reads.

### Single-cell data analyses

Sequencing data were processed with Cell Ranger, including conversion to FASTQ files, alignment to the human reference genome, and generation of gene-barcode count matrices. Downstream analysis was performed in R using the Seurat package. Cells were filtered based on standard quality-control metrics, normalized, and integrated across samples. Hashtag oligo libraries were used to demultiplex donors, pMHC barcodes were annotated using iTRAP2^15^ and doublets were removed prior to analysis. Dimensionality reduction, clustering, and differential expression analysis were carried out using established workflows within Seurat.

### Targeted TCR sequencing of biopsies

Total RNA was isolated from synovial biopsies stored in RNAlater (Thermo Fisher Scientific) using the RNeasy Mini Kit (Qiagen), according to the manufacturer’s instructions. Extracted RNA was subsequently used for TCR library preparation with the SMARTer Human TCR a/b Profiling Kit v2 (Takara Bio), following the manufacturer’s protocol. Briefly, cDNA synthesis was performed using template-switching technology, followed by amplification of full-length TCR α and β chains. PCR products were purified and quantified before sequencing. Purified libraries were submitted to Novogene for paired-end sequencing on an Illumina platform.

Raw sequencing data were processed with the Cogent NGS Immune Profiler Software (CogentIP, Takara). Downstream analyses, including clonotype frequency distribution, integration with single-cell data, and visualization, were carried out in R using the Seurat package.

### Antigen-Induced Marker (AIM) assay with peptide-pulsed CD3^−^ PBMC targets

AIM assay was performed as described previously^20^. Primary human T cells carrying a recombinant TCR knocked into the TRAC locus by CRISPR-Cas9 were used as effector cells. PBMCs were CD3-depleted by magnetic selection and used as target cells. CD3^−^ PBMCs were pulsed with peptide (1 µM, 0.1% DMSO) for 60 min at 37 °C, washed twice in assay medium, and co-cultured with TCR-knock-in CD8+ T cells at a 1:1 effector-to-target ratio (1×10^5^ cells each) in 96-well round-bottom plates.

Co-cultures were incubated for 18–24 h at 37 °C, 5% CO_2_. Cells were harvested, stained with a viability dye and antibodies targeting CD3, CD8, CD69, and CD137 (BD) and analyzed by flow cytometry (Table S3). Effector cells were identified as live CD3+CD8+ T cells, and AIM+ cells were defined as CD69+CD137+. Frequencies were calculated after subtraction of background activation in irrelevant peptide controls.

## Supporting information

Table_S1_Peptide_Library

Table_S2_Pateint_Data

Table_S3_Antibodies

## Funding

This study has received funding from the European Union’s Horizon 2020 research and innovation program under the Marie Skłodowska-Curie grant agreement No. 899987 and DFF Research Project1 No. 2034-00436B.

## Conflicts of interest

The authors declare no conflict of interest.

## Notes

### Competing Interest Statement

SRH is co-inventor on a patent covering the use of DNA barcode-labeled MHC multimers (WO2015185067 and WO2015188839), which is licensed to Immudex. Remaining authors declare no conflict of interest.

