## Supplementary_Figures for "Autoantigen-specific CD8+ T-cell signature in Rheumatoid Arthritis"

Fig S1 – Barcode screen

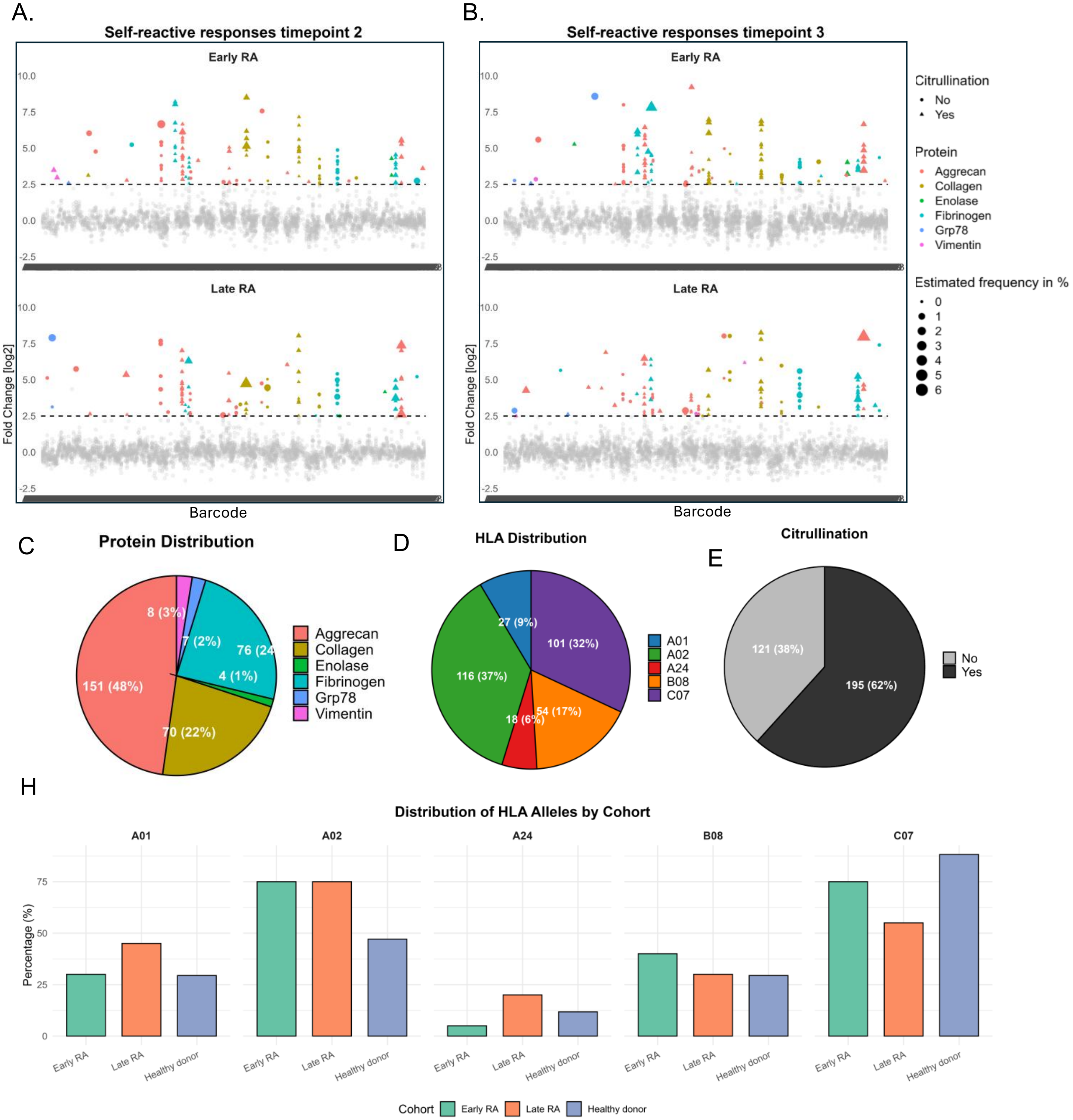


**Figure S1.** Dot plots showing significant (colored) CD8+ T cell responses against RA epitope candidates at timepoint 2 (A) and 3 (B) for early and late RA (triangles = citrullinated peptides, circles = native peptides). Colors refer to proteins and size of the dot indicates the estimated frequency. C) Protein Distribution, (D) HLA Distribution and (E)Citrullination status of all identified 316 RA-specific significant CD8+ T cell responses. H) HLA distribution within HDs and patient cohorts of indicated HLA alleles.

Fig S2– sc- ITRAP2

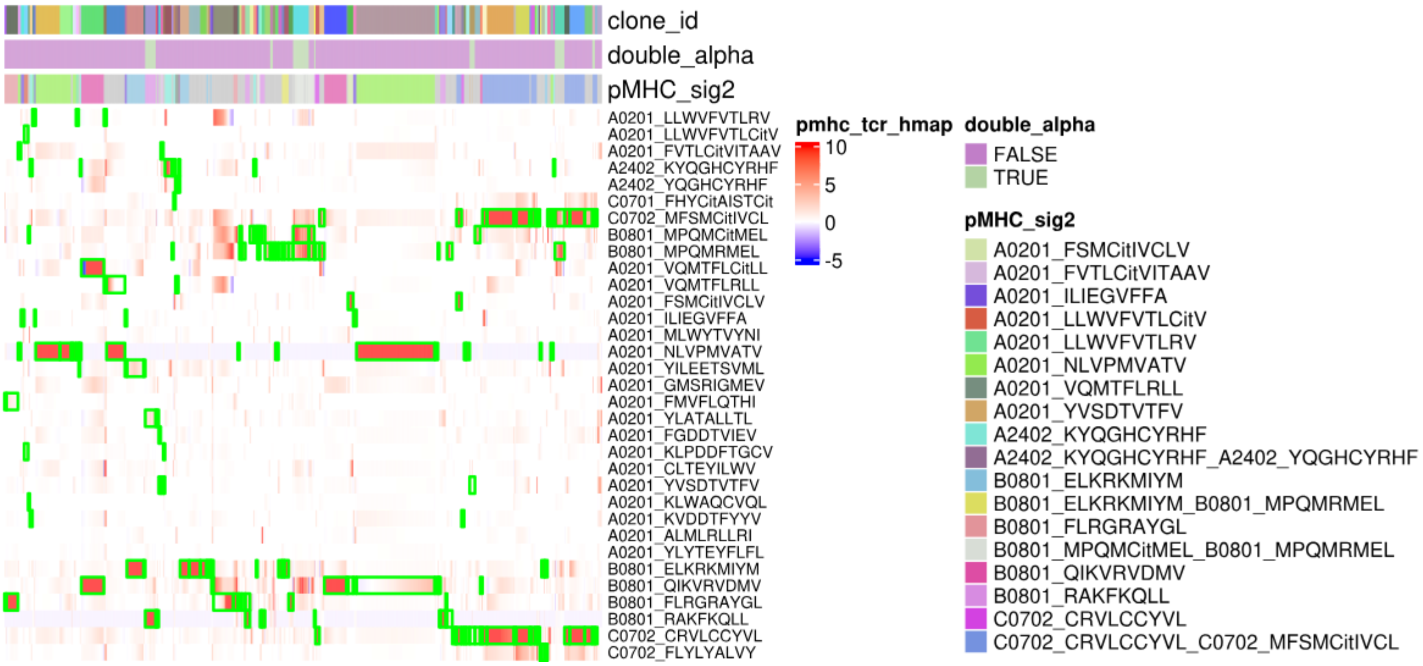


**Figure S2.** Single-cell pMHC barcode–to–T-cell clone annotation performed using iTRAP2, linking antigen specificity to matched TCR clonotypes within the single-cell dataset.

### Fig S3- sc cell type annotations

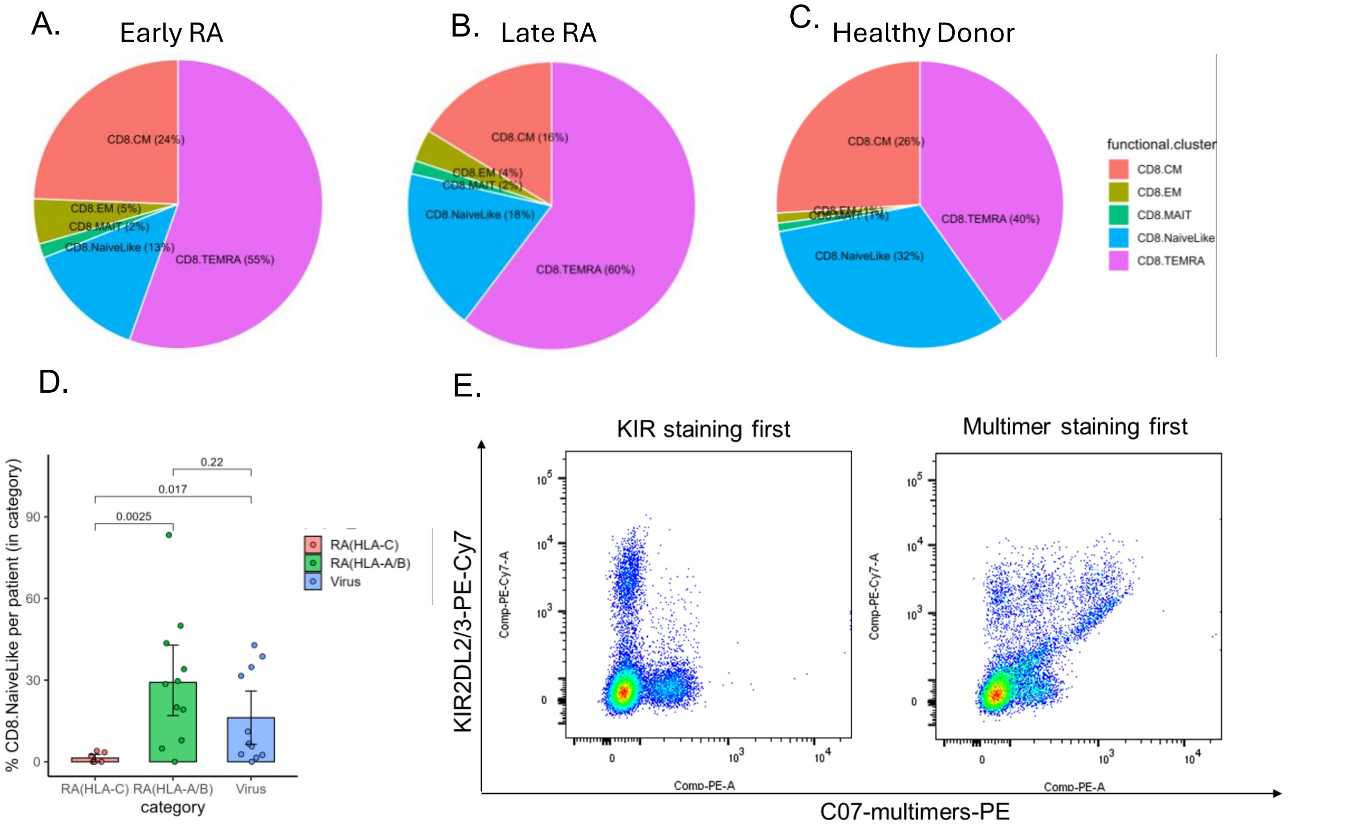


**Figure S3.** Cell type annotations inferred using ProjecTIL for Early RA (A), Late RA (B), and Healthy Donors (C), showing the relative proportions of naïve, TEMRA, TEM, TCM, and MAIT CD8+ T-cell subsets within each cohort. D) Percentage of CD8+ T cells exhibiting a naïve-like transcriptional signature per patient across antigen categories (RA HLA-C–restricted, RA HLA-A/B–restricted, and virus-specific); each dot represents one patient, boxes indicate median and interquartile range, and p-values denote pairwise comparisons between groups (Wilcoxon rank-sum test). E) Flow plots showing the same sample stained with KIR2DL2/3 and C07-multimer under two different conditions: left, KIR staining performed prior to multimer incubation; right, multimer staining performed prior to KIR staining.

### Fig S4- Structural models

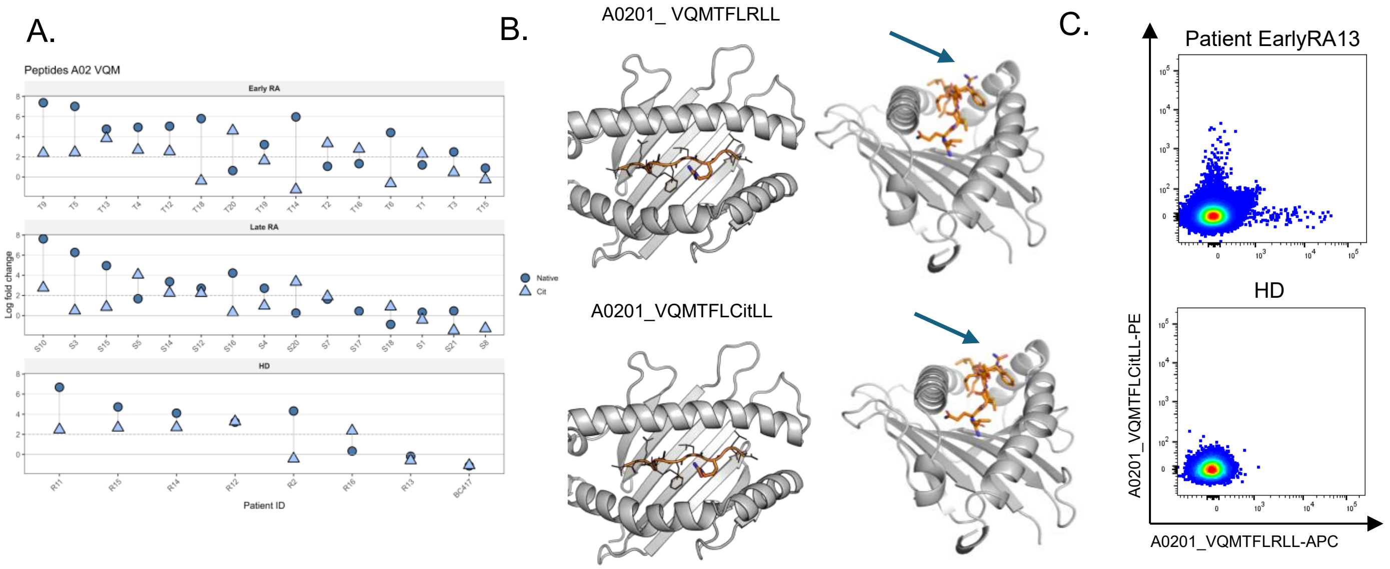


**Figure S4. .** A) pMHC barcode screen data showing responses of HLA-A*02+ patients against RA-specific cit/native A*02-restricted peptide pair VQMTFLR/CitLL. B) Ribbon diagram of the HLA-A*02 monomer (grey) with the bound cit/native peptide ligand depicted (arrow indicates site of citrullination). C) Example FACS plot of tetramer staining with the cit/native peptide pair in patient EarlyRA13 vs HD.

### Fig S5– TCR clonotypes

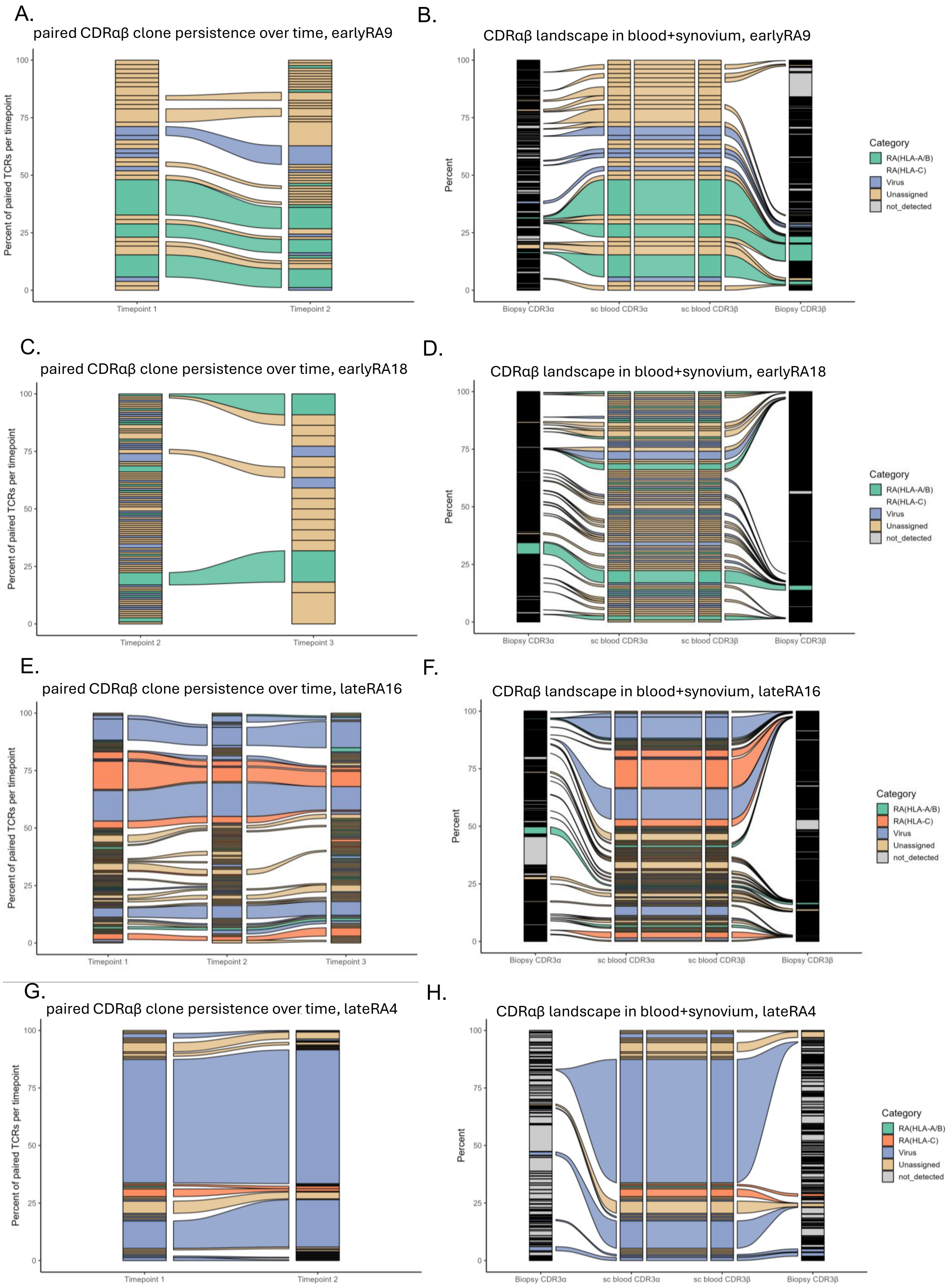


**Figure S5.** A, C, E, G) Alluvial plots showing the persistence and dynamics of TCR clonotypes (paired alpha and beta) in indicated patients across timepoints 1-3. The width of each stream represents the relative frequency of a clonotype, colors correspond to category. B,D,F,H) Alluvial representation of TCR clonotypes shared between blood and synovial tissue compartments. *only one timepoint for patient LateRA9, data not shown.

Fig S6– AIM assay

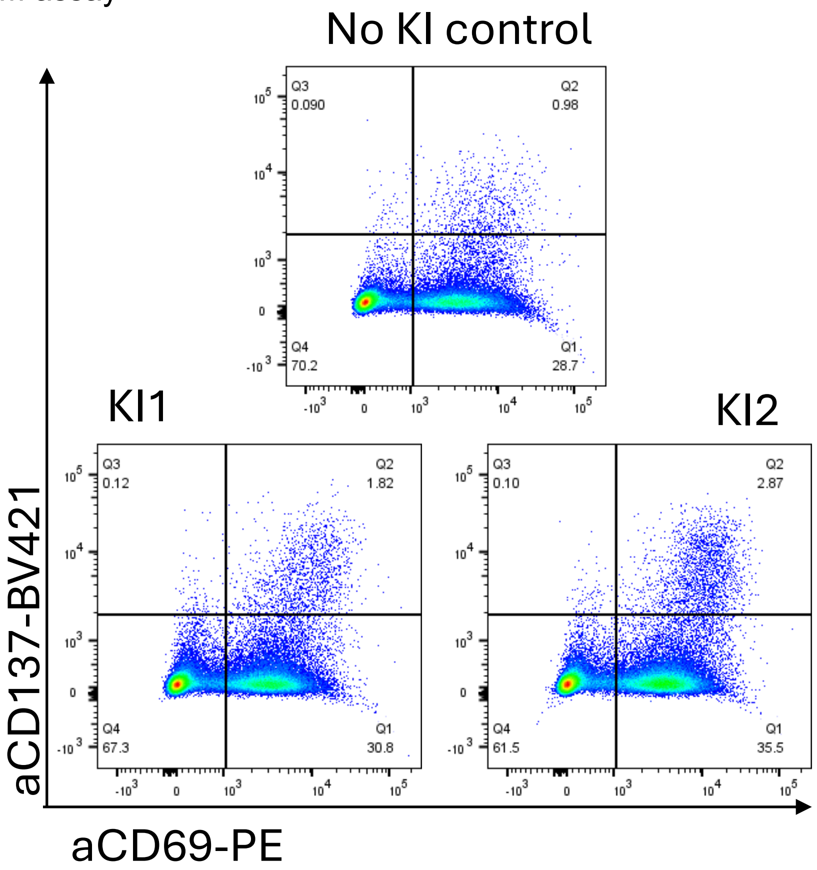


**Figure S6.** Activation-induced marker (AIM) assay measuring CD69 (PE) and CD137 (BV421) expression following co-culture with peptide-pulsed APCs. Upper panel: CD8+ T cells without TCR knock-in (No KI control) stimulated with peptide-pulsed APCs. Lower panels: CD8+ T cells engineered with C07-specific knock-in TCRs (KI1 and KI2) stimulated with the same peptide-pulsed APCs.
